# Adipose tissue microvascular endothelial cells form a tight vascular barrier that selectively transcytoses fatty acid tracers

**DOI:** 10.1101/2024.03.19.585709

**Authors:** Ruby Schipper, Anna Ioannidou, Alice Maestri, Fabiana Baganha, Olivera Werngren, Peder S. Olofsson, Stephen G. Malin, Rachel M. Fisher, Carolina E. Hagberg

**Affiliations:** Division of Cardiovascular Medicine, Department of Medicine Solna, Karolinska Institutet, Stockholm, Sweden; Center for Molecular Medicine, Karolinska Institutet, Stockholm, Sweden

## Abstract

In organs with continuous, non-leaky capillaries like white adipose tissue and the heart, microvascular endothelial cells (ECs) serve as a vital barrier, facilitating nutrient delivery to underlying tissues. While capillary heterogeneity between organs is well established, how these vascular layers have adapted their key functions – such as fatty acid transport – to their respective organs remains unclear, largely due to the lack of organotypic endothelial model systems. Here we demonstrate that the vascular barrier in white adipose tissue, a crucial organ for whole-body fatty acid handling, exhibits comparable impermeability to that of heart and muscle. To investigate if the adipose endothelium possesses tissue-specific functions for facilitating fatty acid transport, we developed an *in vitro* dual tracing-system that allows simultaneous monitoring of barrier integrity and fatty acid transport dynamics by modifying the classic transwell culture. Using this system, we can show human adipose-derived primary ECs selectively transport fluorescent fatty acid tracers while excluding other tracers like dextrans, a phenomenon not observed in other cultured human ECs. Additionally, our findings reveal EC-type specific responses to various transcytosis inhibitors. Our results underscore the unique characteristics of the adipose endothelium and enhances our understanding of how microvascular permeability and transport dynamics have adapted to their specific organ physiology.

## Introduction

Endothelial cells (ECs) are an essential part of the vasculature, covering all luminal vessel walls with a thin monolayer of barrier-forming cells. This ranges from large arteries and veins, lining the complex structures of bigger vessels, to microvascular capillaries which only consist of a single endothelial cell layer, supported by pericytes and a basement membrane, that separate the blood from the parenchyma (1). The structural phenotype of capillary ECs can further be divided into *continuous, fenestrated*, or *sinusoidal* barriers (2). Continuous ECs, typically found in heart and muscle, form a tight non-leaky endothelial layer held together by tight and adherence junctions that effectively prevents para-cellular transport between cells. Instead, continuous ECs engage in trans-endothelial transport, essentially functioning as gatekeepers for the underlying tissues since they form a physical barrier between plasma and tissues; however, this barrier is extremely dynamic. By selectively regulating vascular transport and permeability, the ECs facilitate the exchange of liquids, oxygen, and nutrients in either an active or passive manner. Importantly, this varies between organs, and capillaries ECs from different organs display great transcriptional heterogeneity coupled to functional differences, often reflecting the functions of the underlying parenchyma (3, 4). Despite these insights, much of the mechanistic work studying microvascular ECs has been done on generic endothelial cell lines, not representing the correct EC phenotype.

Interestingly, the white adipose tissue, which under physiological conditions functions as an important storage site for excess fatty acids and other lipid species, also has a continuous EC barrier (2, 5). This suggests the uptake of fatty acids for storage in the adipose tissue to be highly regulated and require active transport into the tissue. Several tissue-specific fatty acid transport mechanisms have been proposed (6, 7). However, how the adipose tissue endothelial permeability and fatty acid transport capacity compares to other continuous vascular beds, such as that of the heart and muscle, has not been functionally explored using tissue-specific primary ECs. This is in part due to the difficult and laborious work associated with obtaining primary ECs from different vascular beds, especially human ECs, forcing the field to rely heavily on the use of human venous umbilical cord derived ECs (HUVECs) that are more accessible (8). Moreover, the use of immortalized endothelial cell lines may not correctly represent cellular phenotypes due to intense subculturing, immortalization-driven defects, and organ of origin and study of study often not matching (9). The increased commercial availability of human primary ECs has now largely abolished this hinder.

Vascular transport of proteins or nutrients, such as fatty acids, is classically studied using transwell cell culture inserts where the transport of fluorescent or radioactive tracers from the apical (top) to basolateral (bottom) can be followed over time. However, the method has remained crude, and distinguishing between changes in EC transport or permeability can be challenging. This is further highlighted by a recent study finding that successful inhibition of trans-endothelial endocytosis (transcytosis) quickly leads to an opening of the vascular barrier, leading to *higher* instead of lower tracer accumulation in the lower well (10). It therefore remains important to further develop organotypic *in vitro* systems such as the transwell model to allow for studies of functional heterogeneity between different types of primary human ECs. Such advances will increase research translatability and permit the study of species specific vascular transport pathways and molecular mechanisms in more detail.

Here we combine *in vivo* and *in vitro* studies to map the organ-specific characteristics of the white adipose tissue microvasculature. Importantly, we have revised the classic transwell system, making it compatible with the study of primary human ECs and enabling simultaneous monitoring of fatty acid transport and barrier integrity in the same well, allowing for comparative monitoring of organ-specific EC functions. We find mouse adipose ECs to possess equally tight vascular barriers as heart and muscle *in vivo*, while *in vitro* using human primary ECs and our optimized system the adipose microvasculature displays a unique substrate selectivity for transporting fatty acid tracers.

## Materials and Methods

### Animal work

All mice were housed in a specific pathogen-free vivarium at the Karolinska Institute. The light/dark period was 12h/12h, and all mice had ad libitum access to food and water. The Stockholm board for animal ethics approved the experimental protocols. Non-fasted wild-type C57BL/6N mice were injected via the tail vein with 50uL of 30 mg/mL Evans Blue (Sigma, cat. #E2129) dissolved in sterile 0.9% NaCl/PBS and placed back in the cage for 60 minutes until toes and nose appeared blue. The circulatory system was perfused with 0.9% NaCl solution until the liver got clear. Subcutaneous and epididymal white adipose tissue (scWAT and eWAT), liver, heart, and muscle were collected in room temperature PBS. Evans Blue was extracted from tissue pieces for 48 hours in N,N-Dimethylformamide (Sigma, cat. #D4551) at 55°C. The solid fraction was removed by centrifugation and in the liquid phase, containing extracted Evans Blue, absorbance was measured at 500nm, 620nm, and 740nm.

### Cells

Primary microvascular ECs purchased from ScienCell Research Laboratories (Carlsbad, USA) or Lonza (BioNordika, Sweden) were used throughout the study. Human adipose microvascular endothelial cells (hAMECs, cat. #7200) are of subcutaneous origin from healthy donors, the human cardiac microvascular endothelial cells (hCMECs, cat. #6000) are isolated from human fetal heart and the human Hepatic Sinusoidal (hLSECs, Lonza #HLECP1) from liver. Human fetal umbilical vein endothelial cells (HUVECS) were a kind gift by Peder Olofsson. Adipose and cardiac ECs and HUVECS were cultured and expanded in Endothelial Cell Medium supplemented with growth factors, P/S and 5% FBS (ECM, ScienCell, cat. #1001). The cells were used between passages 6 to 9, but hLSECs were not expanded and directly plated in transwells. Culture flasks were coated with 2,5 ug/cm^2^ fibronectin (Merck, Sigma Aldrich, cat. #341631) for optimal cell attachment. Cells were subcultured using 0.02% trypsin in 10 mL Phosphate Buffered Saline (PBS), according to ScienCell’s culturing protocol. None of the cultures or treatments showed signs of cell cytotoxicity using the using the LDH-Cytotoxicity Assay Kit II (Abcam, cat. #ab65393) to analyze 50 ul fresh culture media (not shown), according to manufacturer’s protocol.

### Transwell seeding

Transwell® 24-well polycarbonate membrane cell culture inserts with 0.4 μm pores (Corning, Costar, cat. #CLS3413) were coated overnight with 5 ug/cm^2^ fibronectin (Merck). hAMECs were seeded at a density of 40k cells per well while hCMECs, hLSECs, and HUVECs were seeded at 60k cells per well, unless stated otherwise. For a more humanized *in vitro* model, the ECM for the transwells was supplemented with growth factors, P/S and 2% human serum (VWR/BioWest, cat. #S4190-100). The culture insert’s apical site was supplemented with 300uL and the basolateral site with 800uL media. The cells were grown in the same media for three days, while trans-endothelial electrical resistance (TEER) was monitored daily as an indication for barrier development, before tracer experiments were performed. In case of average raw TEER values below 125 Ω the tracing experiments were postponed for a day.

### Measuring barrier integrity via TEER and fluorescent tracing

Barrier formation and integrity was monitored by measuring TEER using the Millicel-ERS Volthometer (cat. #MERS00002) and Milicell-ERS probe (cat. #MERSSTX01, both from Millipore). TEER was measured from day 1 until the day of experiment and after 1 hour and 4 hours of tracer incubation. In parallel, barrier integrity was assessed by adding non-reactive fluorescently labelled dextran to the upper well and measuring its accumulation in the lower compartment of the transwell over time. Fluorescein isothiocyanate–dextran in the size of 40kDa (FITC-dextran, Sigma, cat. #FS40S) was dissolved in MilliQ at a stock concentration of 25 mg/ml. FITC-dextran was mixed with ECM without serum to create a 50uL/well vehicle of 4 mg/ml concentration, resulting in a final concentration of 0,57 mg/mL in the upper well of the transwell insert. The FITC-dextran was incubated for 4 hours protected from light at 37°C, with sampling timepoints at 15 minutes, 60 minutes, and 4 hours after tracing initiation. For each sampling of 100uL from the lower well, 100uL non-labelled media was added back to the lower well and the loss of fluorescence was accounted for during data analysis. Collected media samples were assayed in a black, clear bottom 96-well plate (Corning, cat. #3904) and a FLUOstar Omega microplate reader used to measure fluorescence at an excitation/emission wavelength of 488/520nm. As a positive control for the maximal free passage of FITC-dextran to the lower well, Transwells without cells were used while maintaining all other culture conditions. As a positive control for endothelial barrier opening cytochalasin B was added to the cells, which inhibits actin filament polymerization and therefore results in an irreversible opening of EC tight junctions. Cytochalasin B at a final concentration of 7 ug/mL (Sigma, cat. #C2743) was added to the upper well and incubated for 15 minutes at 37°C before adding tracers.

### Measuring the transport of fluorescent BODIPY-FA and NBDG tracers

A dual system was developed to trace both 0,57 mg/mL FITC-Dextran and 2 μM of the long-chain fatty acid analogue BODIPY-C12 (558/568) (ThermoFisher, cat. #D3835) in one well. BODIPY-FA was conjugated to a 220 μM of BSA dissolved in ECM media without serum at 37°C for 15 minutes protected from light. To test if changes in each tracer concentration could be accurately measured after combining them both in the same well, a standard curve was made for increasing concentrations of BODIPY-FA diluted in 4 mg/mL FITC-Dextran; and for FITC-Dextran diluted in 2 μM BODIPY-FA, without any cells present. In all tracing experiments sampling was done as described above and the BODIPY-FA and FITC-Dextran fluorescence read at 544/590nm (red fluorescence) and 488/520nm (green fluorescence). Graphs specifying fatty acid transport as *BODIPY-FA/Dextran transport ratios* show BODIPY-FA values divided by Dextran values from the same well. Data is normalized to no cell wells which represent free diffusion of tracers, showing active fatty acid transport with values >1. For the concentration gradient and the time course optimization experiments, only FL-BODIPY-FA (Invitrogen cat. #D3822) was used. Glucose was traced using 2-(N-(7-Nitrobenz-2-oxa-1,3-diazol-4-yl)Amino)-2-Deoxyglucose (2-NBDG, Invitrogen, ThermoFisher, cat. #N13195) combined with 0,57 mg/mL 40kDa tetramethyl-rhodamine Dextran (Invitrogen, cat. #42874). Larger 250kDa FITC-dextran (Sigma, cat. #FD250S) was used in combination with BODIPY-FA (558/568).

### Pharmacological vesicular transport inhibition

For treatments with endocytosis inhibitors, the ECs were pre-treated with clathrin inhibitor Pitstop2 (20 µM, Sigma, cat. # SML1169), caveolae-dependent endocytosis inhibitor Genistein (200 µM, Sigma, cat. #G6649), dynamin-inhibitor Dynasore (50 µM, Sigma, cat. #324410) or macropinocytosis inhibitor Amiloride (100 µM, Sigma, cat. #A7410). Inhibitors were incubated 15 minutes prior to the addition of tracers and the changes in tracer transport or TEER monitored as described above.

### Fluorescence microscopy and BODIPY-positive vesicle quantification

After tracer experiments transwell membranes with ECs were washed thoroughly, fixed using 10% Formalin solution and stained with 20 ug/mL Isolectin GS-IB4 From Griffonia simplicifolia, Alexa Fluor™ 647 Conjugate (IB_4_, ThermoFisher, cat. #I32450) or 25 ug/mL Fluorescein or Rhodamine Lens Culinaris Agglutinin (LCA, Vector Laboratories, cat. #FL-1041/RL-1042), and 0,2 ug/mL DAPI solution diluted in 0,1% PBS-Tween. BODIPY-FA fluorescent vesicles accumulated during tracing experiments while the other dyes were used as counter staining after fixation. Membranes were cut out of the inserts and mounted using ProLong™ Diamond Antifade Mountant (Invitrogen, cat. #P36965) and imaged using Ti2 Nikon confocal microscope with 10x or 20x objective. Quantification of fluorescent BODIPY-positive vesicles was performed using ImageJ. The images were converted to 8-bit and a threshold was established to limit unspecific signal. Cell nuclei and BODIPY-positive puncta were counted applying a cutoff of >100 microns^2 and 0.2> and <20 microns^2, respectively.

### Statistical analysis

The data was analyzed using Student’s T-test, one-way ANOVA with Tukey’s multiple comparison correction (for comparison of timepoints) or two-way ANOVA with Sidak or Tukey’s multiple comparison correction, using the GraphPad Prism software version 9 (GraphPad Software, SanDiego, USA). A p-value of ≤ 0.05 was considered significant. ^*^p<0.05, ^**^p<0.01, ^***^p<0.001, ^****^p<0.0001.

## Results

### Adipose endothelial cells form tight continuous vascular barrier *in vivo* and *in vitro*

To demonstrate that white adipose tissue-derived ECs form a continuous, non-leaky endothelial barrier, similar to that of heart and muscle, we injected wild type C57BL/6 mice intravenously with the dye Evans Blue, and measured dye extravasation in multiple organs one hour later (Fig. 1A). While liver, intestines and kidneys were stained dark blue due to extensive dye leakage through the organs’ permeable microvasculature, the subcutaneous and visceral fat depots remained visibly white (Fig. 1B). This was confirmed by measuring dye accumulation in extracted organs, showing that in mice the vascular barrier of the white adipose tissue is comparably tight as that of heart and muscle (Fig. 1C).

**Figure 1:**
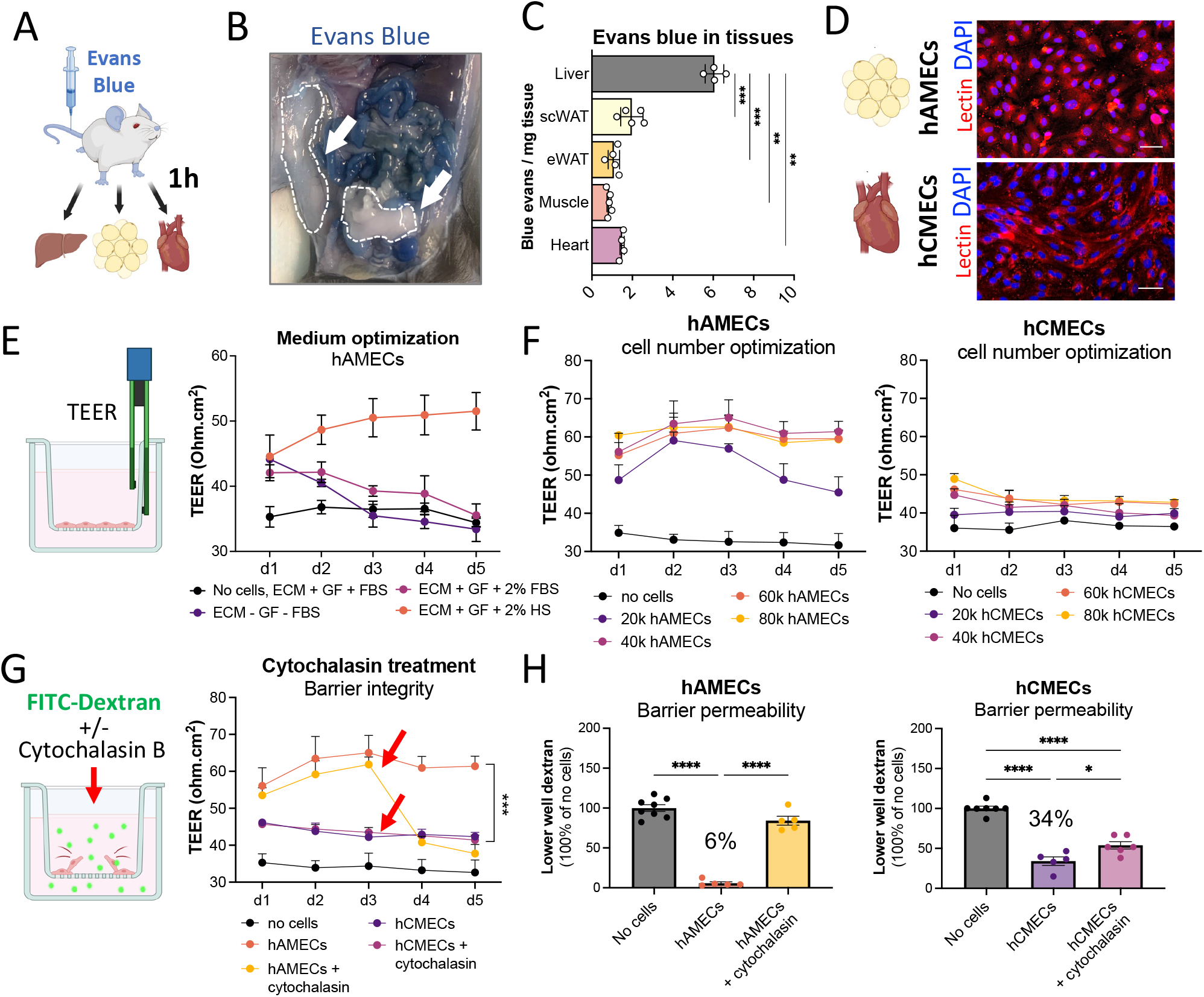
Adipose endothelial cells form tight barrier *in vivo* and *in vitro*. **A)** Schematic of an Evans Blue mouse experiment. **B)** Picture of mouse fat pads and other organs 1h after Evans Blue injection. White arrows and dashed lined indicate subcutaneous and epididymal WAT, respectively. **C)** Quantification of Evans Blue extravasation per organ, normalized to protein. Each datapoints indicated one mouse (n=4-5) from two different experiments. **D)** Confocal images of hAMECs and hCMECs seeded on transwell membranes. Cell membranes are stained with lectin (red) and nuclei with DAPI (blue). Scale bar represents 50 µm. **E)** Schematic of a transwell containing a tight EC monolayer and the electrode used for TEER measurements (left). Measured TEER values (ohm^*^cm^2^) over 5 days for hAMECs plated with different media (right), n=4 wells per data point. FBS= Fetal Bovine Serum, HS=human serum, GF= growth factors. **F)** TEER values over 5 days using different cell seeding densities for hAMECs (left) and hCMECs (right), n=4-8 from 1-2 separate experiments. **G)** Schematic of a transwell containing an EC monolayer disrupted by cytochalasin B while incubated with FITC-Dextran (left). TEER values over 5 days for hAMECs (40k) and hCMECs (60k) incubated with or without cytochalasin B, n=6-10 wells from separate experiments. Red arrows indicate addition of cytochalasin B. **H)** 4 hrs FITC-dextran accumulation in the lower compartment of transwells without cells or plated with hAMECs (left) or hCMECs (right) +/-cytochalasin B treatment. Data is displayed as % lower well accumulation compared to no cells (100%). Each datapoint represents pooled wells (n=4-9) from separate experiments. All data is represented as mean ± SEM. Statistical analyses used are described in the methods section. ^*^p<0.05, ^**^p<0.01, ^***^p<0.001, ^****^p<0.0001.

A tight vascular barrier requires active transport of nutrients and proteins, raising the question how fatty acids and other diet-derived substrates are transported into the adipose tissue. It also asks if differences exist between the microvascular ECs of adipose tissue and heart, two organs with a continuous endothelium but with highly divergent physiological functions and nutritional needs. To probe this in a humanized physiological setting *in vitro*, we set up a transwell culture system compatible with both commercially available primary human adipose tissue-derived microvascular ECs (hAMECs) and human cardiac microvascular ECs (hCMECs). Both EC types formed dense monolayers with prominent cell-cell contacts when plated on fibrinogen-coated transwells (Fig.1D), which could be confirmed by measurements of increased levels of trans-endothelial electrical resistance (TEER) over the vascular layer (Fig.1E). When plated together with human sera instead of commonly used bovine sera, TEER increased continuously from seeding until day three, whereafter it remained stable for at least two more days (Fig. 1E). On the contrary, culturing under serum free (starved) conditions or in the presence of various concentrations of bovine sera did not support the maintenance of an intact hAMEC barrier (Fig. 1E and data not shown). By varying the concentration of plated cells, the optimal seeding density for allowing hAMECs and hCMECs to form a stable monolayer was determined to be 40k and 60k cells per membrane, respectively (Fig. 1F). The difference in number was most due to hAMECs being significantly larger in cell area compared to hCMECs (not shown). When plating fewer or more cells the resistance over the membrane in both cases remained lower, most likely due to too few cells or overgrowth of cells on top of each other, respectively. While both hAMECs and hCMECs formed tight and continuous monolayers with TEER values significantly differing from that of transwells without ECs, the TEER values of hCMECs were on average 10 ohm^*^cm^2^ lower as compared to hAMECs, independent of cell density (Fig. 1F, right). The TEER for hCMECs was not increased by adding more cells or by altering the type of coating of the transwell membranes, suggesting it to be an intrinsic characteristic of the cells and not a culture defect (Fig. 1F and data not shown).

To better understand the differences in TEER between hAMECs and hCMECs we studied their endothelial barrier integrity in more detail. Firstly, we devised a positive control for barrier opening by treating the ECs with the actin polymerization inhibitor cytochalasin B, which prevents actin-dependent endocytosis and thereby causes loosening of the tight junctions that uphold the vascular barrier (Fig. 1G). Treating ECs with cytochalasin B led to a permanent decrease in TEER for hAMECs but not for hCMECs, again suggesting hCMECs form a less tight vascular barrier *in vitro*. To test barrier permeability more thoroughly, we implemented a second method, measuring the passage of a 40 kDa large FITC-labelled dextran over the cellular layer over time. Dextrans are commonly used in permeability experiments as they are not actively transported by ECs, and therefore their passage across the EC layer is used as a measure of either barrier leakage or non-selective transcytosis, depending on if it is transported between cells (leakage) or through them. We used transwells without cells as controls for the maximal accumulation of freely diffusible tracers across the transwell and compared them to FITC-dextran accumulation when ECs were present, allowing us to determine to what extent ECs prevented the passage of dextran to the lower well. Importantly, dextran-tracing is complementary but not identical to TEER measurements, as it detects both transient barrier openings and continuous leakage, while TEER shows only momentary changes in barrier integrity. Measuring FITC-dextran over 4 hours showed that hAMECs efficiently prevented 94% of all dextran diffusion to the lower well, which was reverted by opening the vascular barrier with cytochalasin B, confirming adipose ECs form a tight and dynamic endothelial monolayer also *in vitro* (Fig. 1H, left). Surprisingly given the low TEER levels of hCMECs, the cardiac ECs also prevented 66% of dextran diffusion to the lower well, which was partially reversed by cytochalasin B treatment (Fig.1H, right). This shows hCMECs also form a tight vascular barrier *in vitro*. Taken together, we can show adipose tissue ECs form a tight vascular barrier both *in vivo* and *in vitro*, excluding non-specific extravasation of Evans Blue dye in mice and FITC-dextran in transwell cultures, and that hAMECs and hCMECs display different barrier characteristics *in vitro*, potentially due to their different organ of origin. Our data also outlines a set of necessary experiments for setting up EC-layer specific transwell cultures and confirming their barrier integrity, demonstrating that TEER measurements are not always enough to evaluate barrier integrity and should be complemented with further in-depth experiments including dextran-tracing.

### Current transwell methods do not capture the full complexity of fatty acid tracing

We next measured fatty acid transport over the hAMEC and hCMEC vascular layers to test if the two EC-types displayed differences also in fatty acid transport dynamics (Fig.2A). To this end we used a fluorescent albumin-coupled long chain fatty acid mimic, BODIPY-FA, which is actively transported over EC layers, as its accumulation in the lower well correlates linearly with tracer levels (Fig.2B). Using a low tracer level (2 μM) we found that hAMECs much more efficiently transport fatty acids over their endothelial layer than dextrans, transporting 37% of the fatty acid tracer in 4 hours (Fig.2C and 1H). Surprisingly, hCMECs were just as efficient as hAMECs in transporting BODIPY-FA, and now completely resistant to cytochalasin B treatment (Fig. 2C, right). We confirmed the resistance of hCMECs to cytochalasin B by continuously measuring TEER during tracer incubations, finding no significant dip in TEER for hCMECs, while hAMECs readily underwent barrier opening accompanied by lower TEER values in response to cytochalasin B treatment (Fig.2D). However, when imaging the hAMEC and hCMEC transwell membranes at the 4-hour timepoint, we found no difference between the EC-types in the number of BODIPY-FA-filled intracellular vesicles per cell, suggesting both ECs use vesicle mediated transcytosis to transport BODIPY-FA over the vascular layer, while hCMECs do so through an actin-(cytochalasin B) independent mechanism (Fig. 2C, 2E). No dextran could be detected in either EC layer despite several attempts, most likely owing to either the dye washing away during cell preparations or no or too low internalization of the dextran (data not shown). Taken together, our results raised the possibility that the low TEER values of hCMECs and their highly efficient transport of FITC-dextran to the lower well, could stem from the cardiac ECs engaging continuous non-selective transcytosis rather than experiencing vascular leakage, something most methods would not be able to differentiate. We conclude that commonly used *in vitro* measurements of vascular permeability and transport, including TEER measurements, actin inhibition and dextran tracing, may be insufficient to capture the full complexity of differences in endothelial transport dynamics between distinct vascular beds, calling for the development of more detailed analyses and methodologies.

**Figure 2:**
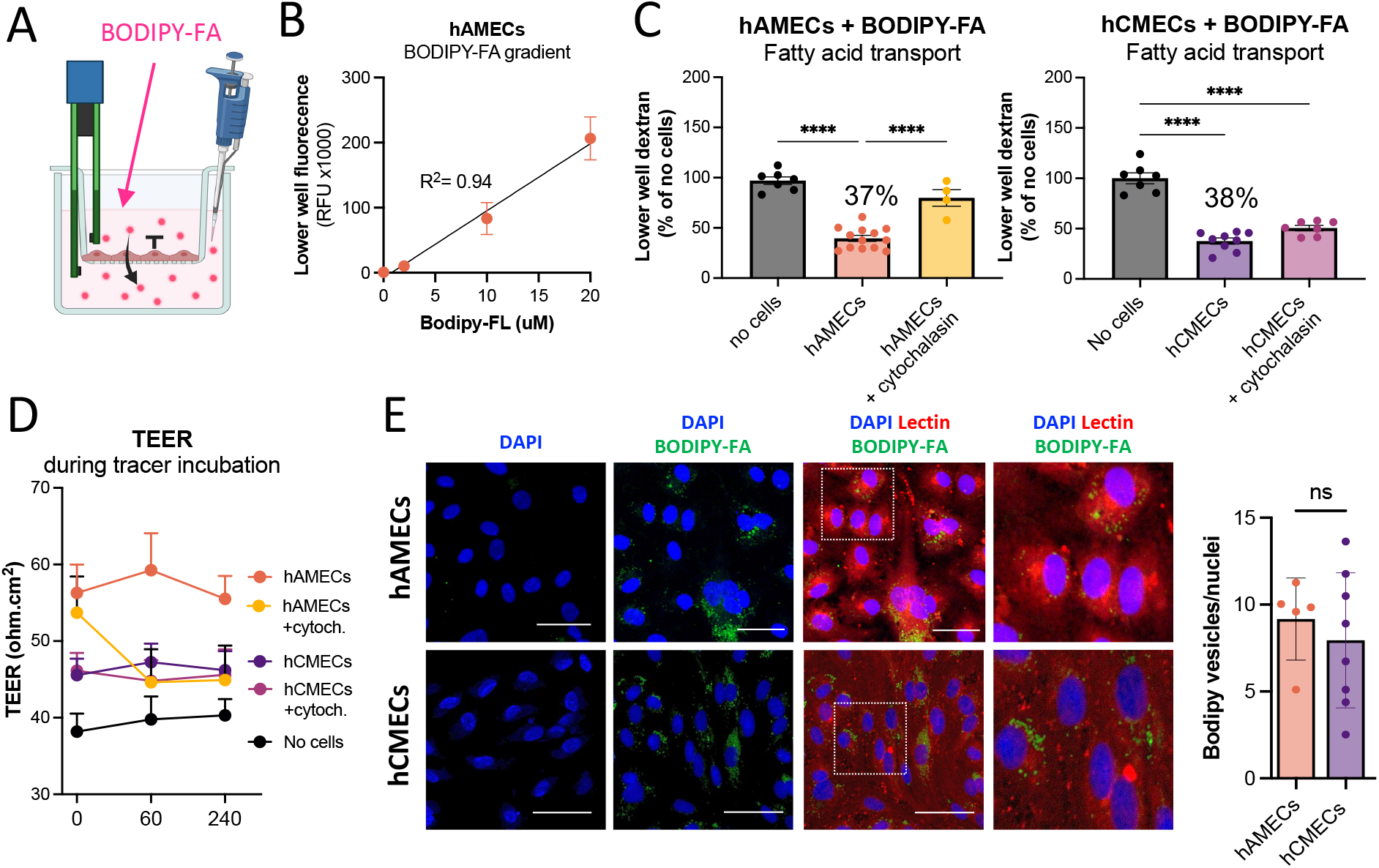
Current methods do not capture the full complexity of fatty acid tracing across transwells. **A)** Schematic of fatty acid tracing in transwell using BODIPY-FA, a TEER electrode, and a pipet for sampling from the lower well. **B)** Accumulated fluorescence in the lower well after incubating transwell-plated hAMECs with increasing concentrations of BODIPY-FA. Each data point represents one well (n=4-5) with fitted linear regression line. **C)** 4 hrs BODIPY-FA accumulation in the lower compartment of transwells without cells or plated with hAMECs (left) or hCMECs (right) +/-cytochalasin B treatment. Data is displayed as % lower well accumulation compared to no cells (100%). Each datapoint represents wells (n=4-12) from separate experiments **D)** TEER measurements for hAMECs or hCMECs +/-cytochalasin B during tracer experiments. **E)** Confocal images of transwell membranes with hAMECs or hCMECs at the 4 hrs timepoint after Bodipy-FA (green) tracing, as indicated. Note the accumulation of clear lipid vesicles. Cells are co-stained with lectin (red), and nuclei with DAPI (blue) before imaging. Scale bar represents 50 µm. Quantification of lipid droplets per cell shown to the right.

### A dual system for fluorescent tracing of both fatty acid transport and barrier integrity

We therefore further adopted the classical transwell system to enable simultaneous assessment of both barrier integrity and fatty acid transport by co-incubating ECs with FITC-dextran and a red-stained (558/568) BODIPY-FA tracer within the same well (Fig. 2A). We normalized the level of accumulated fluorescence in the lower well for each tracer separately when ECs are plated, to tracer accumulation when no ECs were present, which represents the maximal tracer-specific diffusion over time. This allowed us to compare the efficiency of endothelial transport/leakage for each tracer separately within the same well, as well as to each other, normalizing for potential intrinsic differences between the fluorophores (Fig. 3B). Our analytic setup also accounts for differences in EC membrane permeability and allows accurate comparison between different tracer-substrates, cell types and repeat experiments. To test the compatibility of the two tracers together, we incubated increasing concentrations of each tracer with our concentration of choice of the other, finding the two fluorophores did not crosstalk or interfere with each other and that when used together in the same well, both tracers could be accurately detected independently of each other across a range of concentrations (Fig. 3C). Incubating hAMECs and hCMECs with the BODIPY-FA and FITC-dextran mixture showed that while both EC-types continuously transported BODIPY-FA to the lower well, hAMECs did not transport significant levels of FITC-dextran, while hCMECs transported them to the same extent as BODIPY-FA (Fig. 3D). This suggested hAMECS in contrast to hCMECs engage in selective, substrate-specific fatty acid transcytosis while maintaining high barrier integrity. We confirmed this by showing that hAMECs selectively transcytosed only BODIPY-FA, but not the glucose tracer NBDG or a larger dextran tracer to the lower well (Fig. 3E). Another advantage of the dual tracer system is that it allows the mapping of EC-specific phenotypic features and the comparison between different primary human EC layers. By dividing the well-specific transport of BODIPY-FA to that of FITC-dextran, thereby forming a *fatty acid per dextran transport ratio*, we found hAMECs were unique in preferentially transcytosing BODIPY-FA across the vascular layer, while HUVECs and cardiac hCMECs showed equal accumulation of the two tracers in the lower well across all time points, leading to a BODIPY-FA/dextran ratio close to one (Fig.3F). The same was true for human liver sinusoidal ECs (LSECs), that do not form a continuous EC barrier, and for hAMECs treated cytochalasin B to disrupt their barrier integrity, showing our transwell system was able to detect dynamic chances and differences in barrier integrity (Fig.3F). Taken together, by using our dual fluorescent system, the accumulation of BODIPY-FA could be assessed for each transwell in relation to dextran diffusion in the same well, allowing easy comparison between barrier integrity (assessed via dextrans) and nutrient transport (assessed via BODIPY-FA, NDBG or other tracers), comparison between different EC layers, as well as the identification of outlier transwells with damaged EC layers due to adverse handling. Furthermore, our results underline the highly divergent phenotypes of continuous ECs, potentially depending on their organ of origin. They also show primary human ECs show different, vascular-bed specific characteristics *in vitro*, despite being cultured under identical conditions. It suggests organ-specific endothelial responses are maintained in our system and highlights the importance of using primary organ-specific ECs, when possible, for studies of vascular dynamics.

**Figure 3:**
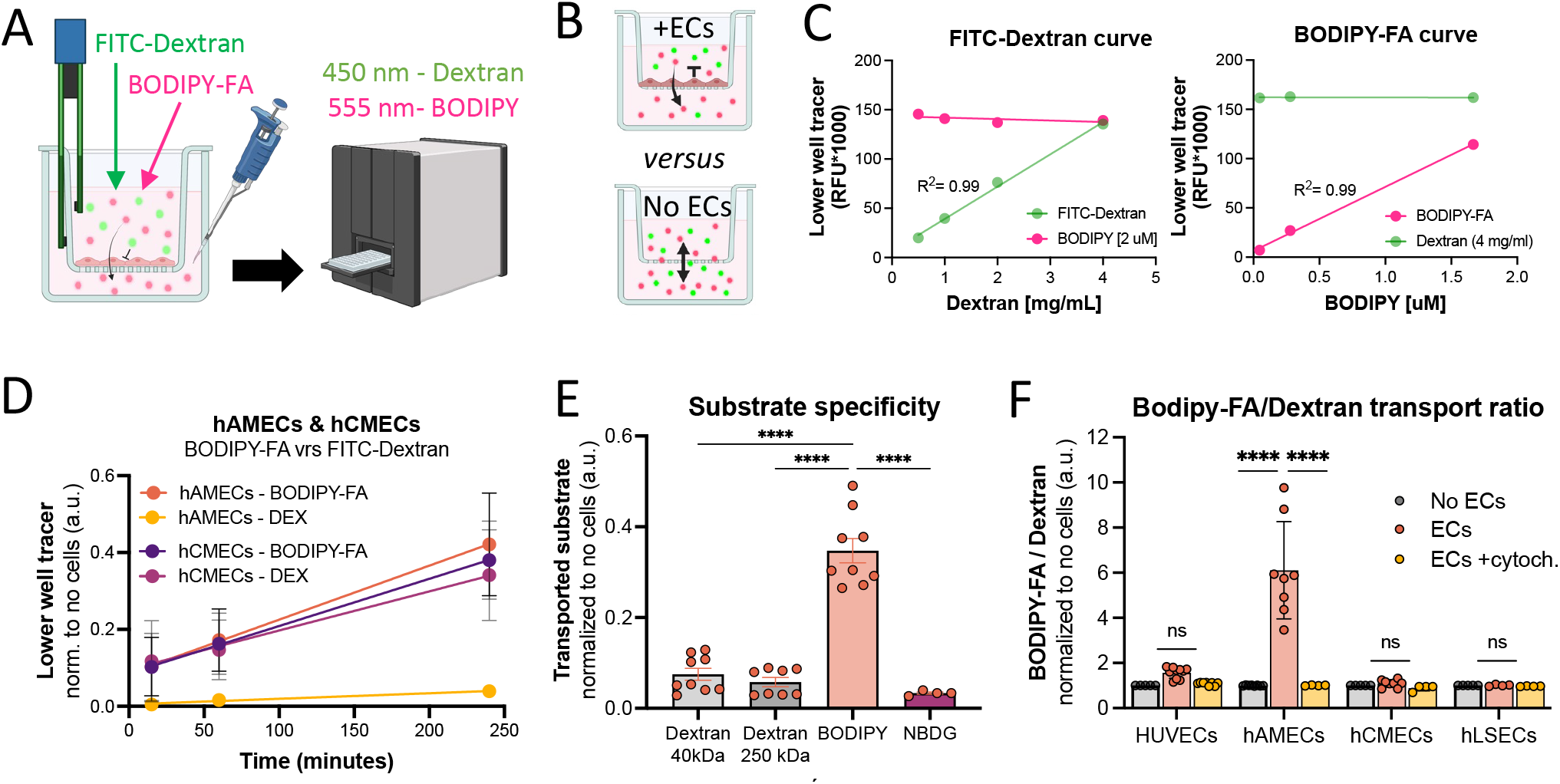
A dual fluorescent tracing system for both fatty acid transport and barrier integrity. **A)** Schematic of an EC-plated transwell incubated with both the BODIPY-FA and FITC-dextran tracers, followed by fluorescent reading at 544/590 nm (red BODIPY-FA fluorescence) and 488/520 nm (green dextran fluorescence) using a plate reader. **B)** Schematic of the analysis, normalizing the level of selective transport over the EC monolayer to that of the maximal versus free diffusion of the tracers to the lower wells in transwells without cells. **C)** Measurements of both dyes while increasing the concentration of FITC-Dextran in a solution of 2 μM BODIPY-FA (left) or increasing BODIPY-FA concentrations in the presence of 4 mg/ml FITC-Dextran (right) with fitted linear regression lines. **D)** Tracer-specific accumulation of FITC-Dextran and BODIPY-FA in the lower well for hAMECs and hCMECs over time. Datapoints represent the average of 4 wells per experiment (n=3-4 experiments), **E)** Transport of different substrates (Dextran 40kDa, Dextran 250kDa, BODIPY-FA, and the glucose tracer NBDG) by hAMECs, showing substrate-preference for BODIPY-FA. Each datapoint represents one well (n=4-9) from experiments. **F)** Fatty acid to dextran transport ratio for hAMECs compared to other primary ECs -/+ cytochalasin B treatment. All data is represented as mean ± SEM from seperate experiments.

### Application of transwell system for comparing different vascular beds

The differential phenotypes of hAMECs and hCMECs let us to ask if the two EC types engage different vesicle-mediated transport pathways, and if identifying these pathways could help to fully establish if hCMECs experience vascular leakage or non-selective transcytosis. We therefore incubated both cell types with a set of small molecule inhibitors targeting each major class of transport vesicles. Caveolae are sensitive to certain tyrosine kinase inhibitors (such as *Genistein*) and to the inhibition of dynamin (using *Dynasore*). Clathrin coated vesicles can be inhibited using *Dynasore* or the more specific inhibitor *Pitstop 2* (PS2), while *Amelioride* often is used to inhibit macropinocytosis. Importantly, all other types of vesicles fully rely on actin polymerization (which is inhibited by cytochalasin B) except clathrin-coated vesicles. Using this set of inhibitors, we could show that while fatty acid transcytosis by the hAMECs showed the typical pattern of drug-sensitivity for caveolae, hCMECs had a very different response, most likely engaging clathrin-vesicles for their fatty acid transport (Fig.4A). The results from using tracers were mirrored by parallel measurement of TEER. Disruption of caveolar dynamics with either *Dynasore* or *Genistein* slightly disrupted the hAMEC’s barriers and thereby decreased their TEER (Fig.4B, left), similar to the effect of cytochalasin (Fig.2D) and to what has previously been reported for primary microvascular endothelial cells of dermal origin (10). In contrast, treatment of hCMECs with clathrin-inhibitors *Dynasore* or *Pitstop 2* instead increased their TEER to levels close to that of hAMECs, with cytochalasin B again showing no effect on TEER. This confirms the two EC layers most likely not only use different trans-endothelial transport mechanisms, but also react differentially to their inhibition, complicating such analyses. It should be noted that all observed effects on tracers and TEER were mild, suggesting the cells may also harbor other parallel transcytosis routs that dampen the effects of the inhibitors.

**Figure 4:**
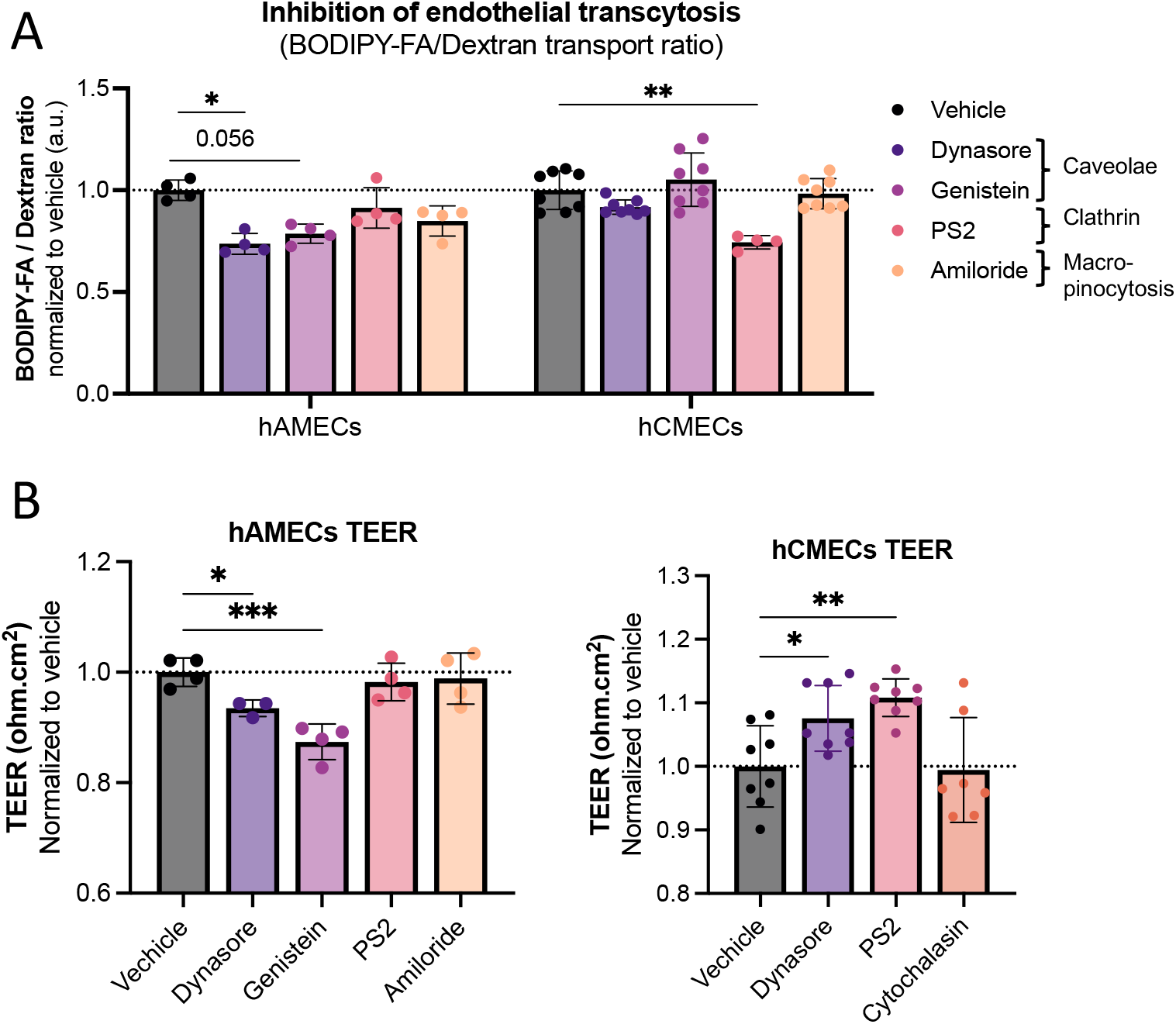
Differential response to endocytosis inhibitors. **A)**Ratio of BODIPY-FA to FITC-dextran accumulation in the lower well of hAMECs (left) and hCMECs (right) after 1 hr incubation with tracers and endocytosis inhibitors targeting different pathways: vehicle (0,1% DMSO), 50µM Dynasore, 200 µM Genistein, 20 µM PitStop2 or 100 µM Amiloride. Each data point represents one well from 2 experiments. **B)**TEER representing barrier integrity 4 hours after endocytosis inhibitor treatments. Each data point represents one well (n=4-8) from a 1-2 experiments. Data represented as normalized to vehicle ± SEM.

Taken together, we show that two types of continuous primary human ECs, derived from adipose tissue and heart, respectively, use very different fatty acid transport mechanisms when cultured *in vitro*. Transport dynamics can be functionally monitored using this dual fluorescent transwell system complemented with TEER measurements, and together with small molecular inhibitors allowing the distinction between non-selective transport and vascular leakage, but care should be taken when interpreting the results from all these methods.

## Discussion

The heterogeneity of ECs among different vessels and tissues has become a well-established phenomenon that requires consideration when studying ECs *in vitro*. While often ignored, the development of novel culture systems that allows the use of tissue-specific ECs is of great importance and will help drive the field forward. New insight will increase our understanding of vessel physiology and allow us to pinpoint mechanisms that become disrupted during disease. Here we adopted the classic transwell system to simultaneously study barrier function and fatty acid transport in a humanized and tissue-specific manner, using human serum and primary human microvascular ECs derived from adipose tissue and heart, respectively. The commercial availability of primary ECs and the range of fluorescent tracers favors widespread adaptation of this technique. We use it to show that human EC-layer specific phenotypes can be preserved in culture and differ greatly between different types of vascular beds. Importantly, our results highlight that many common transwell methodologies do not fully capture the complexity of endothelial dynamics, and more analytical systems and measurement models should therefore be developed. The use of two tracers in one well increases data collection from one single experiment and allows for easy analyses that can be performed in any lab. It also allows for straightforward and transparent exclusion of outliers where the barrier integrity has been damaged due to technical misses. Most importantly, the two ways of measuring barrier integrity allow for parallel internal controls within the system.

While the use of primary ECs poses problems such as their potential dedifferentiation during even low passaging, many of their cell-typic features have been shown to remain *in vitro* (11-13). However more studies are needed to understand how to preserve native organotypic endothelial phenotypes in culture even better. Interestingly, single cell sequencing in mice revealed the transcriptomes of adipose and cardiac ECs to be most similar to each other, yet not identical, as compared to other isolated capillary ECs from mouse (14, 15). Despite both hAMECs and hCMECs being categorized as continuous EC barriers *in vivo*, we show that their molecular transport mechanisms for fatty acids differed when cultured *in vitro*. While hAMECs showed substrate-specific transport of only the BODIPY-FA tracer, hCMECs co-transported BODIPY-FA and dextran non-selectively, most likely within clathrin-coated transport vesicles judging from treatment with endocytotic inhibitors. This is not surprising, as clathrin-dependent transcytosis is considered the main vesicle-mediated uptake pathway for continuous endothelium (16). Inhibition of clathrins reduced hCMEC fatty acid transcytosis and increased their TEER. This strengthens our conclusion that that cultured primary hCMECs do not suffer from poor barrier integrity, as could have been easily judged from our TEER data (Fig.1G), but most likely engage in continuous non-selective transcytosis, mixing the solutes from the apical and basal wells, and thereby “diluting” the measurement of TEER which is sensitive to cell culture composition (17). This hypothesis would also fit with lower TEER values observed for hCMECs during culture day 1-3 despite the use of human sera (Fig.1F).

Based on literature it was instead more surprising that the data presented in Fig.4 suggest hAMECs to use caveolae for fatty acid uptake (16). Both the adipose and heart microvasculature is very rich in caveolae, but these vesicles have by some been suggested to mostly conform a structural storage site for endothelial cholesterol, and not engage in endocytosis. Whereas this is true remains to be confirmed, most notably by knocking out specific vesicle components such as CAV1 for caveolae in our cultures, and/or using specific knockout mice. Because both hAMECs and hCMECs harbor caveolae, and their uncertain role in fatty acid transcytosis, we conclude that the vesicular mechanism mediating substrate specificity for the adipose endothelium remains to be determined, despite the apparent EC-specific pattern of inhibitor-sensitivity to caveolar transport. The involvement of specific fatty acid transporters should also be investigated since these receptors can reside inside budding sites of endocytic vesicles (16, 18, 19). For instance, our system could be utilized in combination with genetic knockdown of the CD36-inhibitor Sulfo-N-succinimidyl oleate (SSO) (15, 20) or molecules that inhibit mitochondrial ATP production (21). The ECs could also be treated with known inducers of organ-specific vascular transcytosis, such as VEGF-B (22) or Angiopoietin II (23). This shows the diversity of applications adaptable to the system, and further strengthens its utility for answering a variety of research questions.

While differences in EC beds remain to some extend visible *in vitro*, we should bear in mind that the lack of supporting cells and parenchyma might affect cellular transport. For instance, although transwells allow for cell polarization towards and apical and basolateral site which facilitates transport, the membrane is more rigid compared to the endothelial basal membrane and is of synthetic origin that could have implications for cell morphology (24, 25). Progress is being made by developing biological membranes such spider silk protein-based membranes that are thinner and more flexible (26, 27). Moreover, ECs are *in vivo* always exposed to flow and sheer stress in circulation (28), another component missing in this transwell system. However, the complexity, costs and expertise needed contribute to the fact that full engineered systems containing these components are still scarce.

In summary, we have revitalized and further optimized the classic transwell system. Our findings show *in vitro* heterogeneity in endothelial fatty acid transport and barrier dynamics in two types of microvascular endothelial cells that form continuous capillaries, which might stem from the reliance on different vesicular transport mechanisms. Mainly highlighting that substrate specificity for fatty acids in adipose ECs compared to ECs from other origin remained. These results were made possible by using a dual tracer incubation of FITC-Dextran and BODIPY-FA together, allowing simultaneous monitoring of EC barrier integrity and fatty acid transport. By only culturing primary ECs and using human serum, the translatability and biological relevance of the results will most likely be improved. Importantly, we show that this system can be manipulated using barrier disrupting agents or transport inhibitors and that it provides an easy-to-use *in vitro* application that is adaptable to a large variety of specific research questions.

## ACKNOWLEDGEMENTS and FUNDING

We thank Anneli Olsson for valuable technical assistance and Volker Lauscke for providing LSECs. CEH was supported by the Swedish Research Council (#2019-02046) and Karolinska Institutet (2-189/2022 and 2020-00893).

## AUTHOR CONTRIBUTION

RS, AI, RMF CH conceptualized and planned the study and RS, AI, FB and OW performed the experiments. RS, AI, AM, FB and CH analysed the data. OW provided technical assistance. Animal work was done by FB and CEH in collaboration with PSO and SGM. RS wrote the first version of the manuscript and CH and RS edited it together.

